# Expression and function of voltage gated proton channels (H_V_1) in MDA-MB-231 cells

**DOI:** 10.1101/2019.12.23.886945

**Authors:** Dan J. Bare, Vladimir V. Cherny, Thomas E. DeCoursey, Abde M. Abukhdeir, Deri Morgan

## Abstract

Expression of the voltage gated proton channel (H_V_1) as identified by immunocytochemistry has been previously reported in breast cancer tissue. Increased expression of H_V_1 was correlated with poor prognosis and decreases overall and disease-free survival but the mechanism of its involvement in the disease is unknown. Here we present electrophysiological recordings of H_V_1 channel activity thus, demonstrating their presence and functional properties in the plasma membrane of a breast cancer cell line, MDA-MB-231. With western blotting we also identify significant levels of H_V_1 expression in 3 out of 8 “triple negative” breast cancer cell lines (estrogen, progesterone, and HER2 receptor expression negative). We examine the function of H_V_1 in breast cancer using MDA-MB-231 cells as a model by suppressing the expression of H_V_1 using shRNA (“knock-down; “KD”) and by eliminating H_V_1 using CRISPR/Cas9 gene editing (“knock-out”; “KO”). However, these two approaches produced different effects. Knock-down of H_V_1 using shRNA resulted in slower cell migration in a scratch assay and a significant reduction in H_2_O_2_ release. In contrast, H_V_1 KO cells do not show reduction in migration or H_2_O_2_ release. H_V_1 KO but not knock-down cells showed an increased glycolytic rate with an accompanied increase in p-AKT (Phospho-AKT, Ser473) activity. The expression of CD171/LCAM-1, an adhesion molecule and prognostic indicator for breast cancer, was reduced in the absence of H_V_1 expression. When we compared MDA-MB-231 xenograft growth rates in immunocompromised mice, tumors from H_V_1 KO cells grew less in mass with lower staining for the Ki-67 maker for cell proliferation rate. Therefore, deletion of H_V_1 expression in MDA-MB-231 cells limits tumor growth rate. The limited growth thus, appears to be independent of oxidant production by NADPH oxidase molecules and be determined through cell adhesion activity. While H_V_1 KO results in cell mechanisms different from KD, both implicate H_V_1-mediated pathways for control of tumor growth in the MDA-MB-231 cell line.

## Introduction

The voltage gated proton channel (H_V_1), part of the superfamily of voltage-gated membrane proteins, is a membrane bound 273 amino acid protein that forms a pH- and voltage-gated ion channel that conducts protons [1, 2]. It forms a dimer in the membrane in which each monomer has four membrane spanning subunits (S1-S4) and each monomer has its own proton-conducting pathway [3–5]. When the channel opens it is perfectly selective for protons [6–9]. The channel senses the pH gradient across the cell membrane and opens when the electrochemical gradient for H^+^ is outward, resulting in acid extrusion that raises pH of the cytosol [10]. As it extrudes H^+^, it is electrogenic and causes hyperpolarization of cellular membranes. During the respiratory burst of phagocytes, it facilitates and sustains the activity of the enzyme NADPH oxidase by compensating for both pH and membrane potential changes that would otherwise inhibit the enzyme’s function [11–14]. A close functional relationship with the NADPH oxidase is also seen in B cell receptor signaling [15] and in pathophysiological states in ischemic stroke where NADPH oxidase in microglia contributes to bystander injury facilitated by H_V_1 [15, 16]. Important physiological effects of H_V_1 on cytosolic pH have also been demonstrated during histamine release by human basophils [17, 18] and in sperm where it contributes to capacitation and motility [17, 18]. In mammals, only a single gene codes for H_V_1 although the channel protein may be truncated in sperm and in chronic lymphocytic leukemia through post-translational processing [19, 20]

H_V_1 protein expression was reported in certain types of breast cancer cells and inhibition of the channel expression by siRNA or shRNA was found to reduce migration, lower the release of matrix metalloproteinase enzymes and produce smaller tumors in a mouse model [21]. A follow-up clinical study showed a correlation between the expression levels of H_V_1 and worse prognosis, lower recurrence-free survival and poor disease scores [22]. However, the mechanisms by which H_V_1 regulate breast cancer cell growth are not known.

One hallmark of cancers is the disordered metabolism seen in a large number of tumors that upregulate the uptake and use of glucose as a source of energy [23, 24]. Many cancer cells utilize glycolysis as the predominant energy pathway, causing local lactic acidosis either due to the tumor microenvironment being hypoxic and requiring anaerobic fuel sources or because the cells have become reprogrammed to utilize glycolysis even in the presence of adequate oxygen (the Warburg effect) [25, 26]. The increased glycolysis and resulting acidosis of the surrounding tissue causes environmental pressure in both cancer cells and the surrounding tissue. Cancer cells adapt to this change by upregulating a number of proton transporters including H^+^-ATPases, Na^+^/H^+^ antiport, carbonic anhydrase, bicarbonate transporters and monocarboxylate transporters resulting in a cytosolic pH (pHi) that is higher (pHi = 7.2-7.4) than the surrounding milieu (pH_o_ ~6.9) [27–29].

The presence of H_V_1 in breast cancer cells and the manner of its function runs contrary to the action of pumps and transporters commonly upregulated in cancer as it only moves protons “downhill” not against the pH gradient required to invert the pH gradient. If we can establish its function we can draw new ideas about how cancer cells control their growth. Here we present further evidence that the proton channel H_V_1 contributes to in-vivo cancer cell tumor growth through mechanisms that are independent of reactive oxygen species (ROS) production.

## Methods

### Cell lines and cell culture

Cells were obtained either from ATCC (American Type Culture Collection, US; HEK297, MDA-MB-231, MCF-7, MDA-MB-468, MDA-MB-436, SKBR3, BT-474, MCF-10A, BT-20, BT-549, Hs578t and MCF-10A) or from Asterand Bioscience (Sum-159PT, Sum-229PE). Cells were grown in a Forma Scientific water-jacketed incubator at 37 °C, 5% CO_2_. DMEM media supplemented with 10% FBS, GlutaMAX, and 25 mM HEPES (Invitrogen, USA) was used for MDA-MB-231, MCF-7, Hs578t, BT-20, MDA-MB-436, MDA-MB-468, Hs578t cell lines. Ham’s F12 with 10% fetal bovine serum, 10 mM HEPES, 1 μg/ml Hydrocortisone and 5 μg/ml bovine insulin media was used to grow SUM-229PE and Sum-159PT. DMEM/F12 with 5% fetal horse serum, 2 ng/ml EGF, 0.5 mg/ml hydrocortisone, 100 ng/ml cholera toxin, and 10 μg/ml insulin was used for MCF-10A cells. All media was supplemented with penicillin/streptomycin at 1000/100 units/ml (Invitrogen).

Cells lines were utilized for fewer than 20 passes. All extracts and transfections from cells were taken from sub-confluent cells in their logarithmic phase of growth unless stated.

### Electrophysiology

Cells were grown to ~80% confluence in 35 mm cultures dishes at 37°C in 5% CO_2_. To detach the cells media was removed and trypsin (0.25 % in EGTA; GIBCO, US) added for 1-2 min. Adding media stopped the trypsinization, then cells were centrifuged for 4 min at 200 x g and resuspended in fresh media. The cells were plated onto glass cover slips at low density for patch clamp recording.

Micropipettes were pulled using a Flaming Brown automatic pipette puller (Sutter Instruments, San Rafael, CA) from Custom 8520 Patch Glass (equivalent to Corning 7052 glass; Harvard Apparatus, Holliston, MA), coated with Sylgard 184 (Dow Corning Corp., Midland, MI), and heat polished to a tip resistance ranging typically 3-10 MΩ with highly buffered tetramethylammonium^+^, TMA^+^, containing pipette solutions. Electrical contact with the pipette solution was achieved by a thin sintered Ag-AgCl pellet (In Vivo Metric Systems, Healdsburg, CA) attached to a Teflon-encased silver wire, or simply a chlorided silver wire. A reference electrode made from a Ag-AgCl pellet was connected to the bath through an agar bridge made with Ringer’s solution. The current signal from the patch clamp (EPC-9 from HEKA Instruments Inc., Holliston, MA, or Axopatch 200B from Axon Instruments, Foster City, CA) was recorded and analyzed using Pulse and PulseFit software (HEKA), or P-CLAMP software supplemented by Sigmaplot (SPSS Inc., Chicago, IL). Seals were formed with Ringer’s solution (in mM: 160 NaCl, 4.5 KCl, 2 CaCl_2_, 1 MgCl_2_, 5 HEPES, pH 7.4) in the bath, and the potential zeroed after the pipette was in contact with the cell. Current records are displayed without correction for liquid junction potentials.

The whole-cell configuration of the patch-clamp technique was used. Bath and pipette solutions were used interchangeably. They contained (in mM) 2 MgCl_2_, 1 EGTA, 80-100 buffer, 75-120 TMA^+^ CH_3_SO_3_^-^ (adjusted to bring the osmolality to ~300 mOsm), and were titrated using TMAOH. Buffers with p*K_a_* near the desired pH were used: Homopipes for pH 4.5-5.0, MES for pH 5.5-6.0, BisTris for pH 6.5, BES for pH 7.0, HEPES for pH 7.5, Tricine for pH 8.0, and CHES for pH 9.0. Experiments were done at room temperature (~20-25°C). Current records are shown without leak correction.

Reversal potentials (*V*_rev_) in most cases were determined from the direction and amplitude of tail current relaxation over a range of voltages, following a prepulse that activated the proton conductance, *g*_H_. Currents were fitted with a single exponential to obtain the activation time constant (*τ*_act_) and the fitted curve was extrapolated to infinite time to obtain the “steady-state” current amplitude (*I*_H_), from which the *g*_H_ was calculated as *g*_H_ = *I*_H_/(*V*-*V*_rev_). Thus we assume that the time dependent component is due to H^+^ current and time independent current represents leak. Because of the strong voltage dependence of activation kinetics, we frequently applied longer pulses near threshold voltages, and shorter pulses for large depolarizations in order to resolve kinetics and avoid proton depletion associated with large H^+^ flux. The voltage at which *g*_H_ was 10% of *g*_H,max_ (*V*_*g*H,max/10_) was determined after defining *g*_H,max_ as the largest *g*_H_ measured.

### Membrane protein isolation and western blotting

Cells growing exponentially in culture were scraped using a cell scraper (Midwest Scientific) and pelleted by centrifugation at 200 x g for 4 min at 4 °C. Cells were resuspended in PBS and washed twice by centrifugation and resuspension. Cell pellets were resuspended in ice cold homogenization buffer (140 mM Tris-HCl, EGTA 10 mM) supplemented with protease inhibitors (HALT, Invitrogen, US). The cells were lysed by being drawn through a 27-gauge needle 15-20 times using a 1 ml syringe while being kept at 4 °C. The resulting cell lysate was centrifuged at 16,000 x g for 30 min. The supernatant was discarded and the resulting membrane pellet was resuspended in fresh buffer with protease inhibitors. The protein concentration of the membrane samples was determined using a micro BSA protein assay kit (Thermo Scientific, USA) and the samples were frozen at −80 °C until further use. Cell membrane samples were rapidly thawed and 250 μl of Laemmli sample buffer (Bio-Rad Labs, Hercules, CA) was added. This solution was boiled for 10 min at 100 °C and protein isolates were resolved using SDS-PAGE using 4–12 % Bis–Tris NuPAGE gels in MES running buffer (Invitrogen, Grand Island, NY) following the manufacturer’s protocol. The proteins were transferred using Invitrogen Xcell II blotting apparatus to a PVDF membrane (Invitrogen, Grand Island, NY). Following transfer, the membranes were blocked in 5% w/v blotting-grade blocker (Bio-Rad Labs, Hercules, CA) in tris(hydroxymethyl)aminomethane (TRIS)-buffered saline supplemented with 0.1% Tween-20 (Sigma, Saint Louis, MO) for 1 h. Membranes were probed with primary antibodies to H_V_1 (kind gift from Melania Capasso, Deutsches Zentrum für Neurodegenerative Erkrankungen in der Helmholtz-Gemeinschaf), NHE-1, AKT, phosphoAKT_s473 and Na^+^/K^+^ ATPase (Cell Signaling, Beverly, MA) Cas9 (TakaraBio, Mountain View, CA) followed by incubation with an anti-rabbit secondary antibody conjugated to horseradish peroxidase (Cell Signaling, Beverly, MA). Protein bands were visualized using enhanced chemiluminescent reagent (Perkin-Elmer, Waltham, MA) and HyBlot-CL autoradiography film (Harvard Bioscience, Inc.). Densitometry was performed using ImageJ analysis software (NIH).

### Immunostaining

MDA-MB-231 cells were harvested by brief trypsinization followed by plating onto 12 mm round coverslips (thickness, #1.5) coated with poly-D-lysine. Following overnight attachment in medium, the cells were rapidly washed with warm PBS and then fixed with the addition of 4% paraformaldehyde in 200 mM phosphate buffer (pH 7.4) for 30 min. Cells were permeabilized with 0.3% Triton X-100 in phosphate-buffered saline. After Triton X-100 washout, cells were blocked overnight in buffer containing 10% goat serum. The expression of the proton channel was detected by incubation with a rabbit polyclonal primary antibody raised against human HVCN1 (gift from Melania Capasso, Deutsches Zentrum für Neurodegenerative Erkrankungen in der Helmholtz-Gemeinschaf) used at a 1:100 dilution for 3.0 h. Following a washout period, coverslips were incubated with an Alexa-fluor 488-conjugated anti-rabbit secondary antibody. After a secondary antibody washout period, the cells were mounted in DAPI Fluoromount-G (Southern Biotech, Birmingham, Al) for nuclear co-labeling. All coverslips were evaluated with an inverted Leica microscope and confocal microscopy. Confocal images (1.0 μm) were acquired from the cell surface to the bottom for determination of sections through the center of the nucleus.

### Creation of luciferase expressing, HVCN1 knockdown and knockout cells

Lentiviral HVCN1 knockdown mission shRNA plasmids (SHCLNG MISSION shRNA Bacterial Clone Olig# TRCN0000165728, TRCN0000161821, TRCN0000162585) were obtained from Sigma (Cambridge, MA, USA). The lentiviral packaging and envelope vectors psPAX2 and pMD2.G (12259 12260), pLKO-control shRNA [30] (Addgene plasmid 1864 deposited by Dr. David M. Sabatini), LentiCas-9BLAST [31]; were obtained from Addgene plasmid repository. Luciferase (red) lentiviral vector (Luciola Italica) (pLenti-II-CMV-Luc-IRES-GFP) was obtained from ABM (Richmond, BC, Canada). The CRISPR guide sequence (gRNA) was designed using crispr.mit.edu and the sequence of HVNC1 from the UCSC Genome Browser (NM_0322369). The gRNA guide sequence (TTAAGGCACTTCACGGTCGT) was ligated into a pLentiGuide-Puro or pLentiGuide-Neo plasmid (Genscript). Lentiviral supernatants for the expression of shRNAs, Cas9 plasmids, luciferase plasmids and gRNA plasmids were generated from 293FT cells using psPAX2 and pMD2G packaging and envelope vectors or 2^nd^ generation packaging mix from ABM (Richmond, BC, Canada). HEK293FT cells were grown to 40 % confluence in 75 ml tissue culture flasks. Media was removed and replaced with DMEM, 10 % FBS without antibiotics. Fugene 6 (PROMEGA, Madison, WI) or Lentifectin (ABM, Richmond, BC, Canada) was used to transfect the psPAX2, pMD2.G and expression plasmid in a ratio of 2:1:3 concentration or using the manufacturers protocol. After 48 hours the virus supernatant was collected and filtered through a 0.44 mm syringe filter and then concentrated using Lenti-X concentrator (Takara biological, Mountain View, CA) following the manufacturer’s protocol and resuspended in the appropriate media. To transduce cells with virus, cells were grown in 6 well cell culture plates; the media removed and replaced with 1 ml of the virus media added to the wells supplemented with polybrene. After 4-24 hours of incubation the media was replaced with fresh media and the cells were left to recover overnight. Transformed cells were then trypsinized and replated in media containing a selection chemical (20 μg/ml blasticidin, 2 μg/ml puromycin or 600 μg/ml neomycin). After the colony stabilized from the selection challenge, 200 μl of a cell suspension was plated at a seed density of 1 cell/ 200 μl into each well of a 96 well plate. Single cell colonies formed in the wells, were isolated and grown up to confluence in 6 well tissue culture flasks. Protein isolates were made from successful colonies and assayed for the expression of various proteins by western blot. Primers flanking the Cas9-mediated cut site were used to amplify this region for subsequent sequencing to determine the change to the sequence of the gene. Putative off-targeting sites that were predicted by crispr.mit.edu were also sequenced to determine if off-targeting of the Cas9 protein occurred.

### In-vivo measurement of xenograft growth

32 two-month-old NOD.Cg-Prkdc<scid> female mice (Jackson Labs, Sacramento, CA) were housed in groups of 6 mice under standard 12 hour light/dark cycles with food and water ad libitum. The number of mice required was determined using power analysis to achieve 0.80 power (at alpha = 0.05) to detect differences of 20% or greater (ANOVA statistical test; validated using StatMate2 [GraphPad Software, Inc., La Jolla, CA]). Based on preliminary and published data we estimated a minimum of 12 mice per experimental and control groups for these experiments. All procedures were approved by Rush University Medical Center Institutional Animal Care and Use Committee (IACUC) in accordance with the NIH Guide for the Care and Use of Laboratory Animals. MDA-MB-231 cells transfected with luciferase gene without any genetic modifications (WT-luc), stably transfected with Cas9 (Cas9-luc) or with HVCN1 genetically deleted (4a-luc) were resuspended at 1×10^6^ cells/100 μl in 50% DMEM/ 50% Matrigel Matrix High Concentration (HC) (Corning, Tewksbury, MA, USA) and kept on ice. Mice were randomly assigned to groups and deeply anesthetized using 3 % isoflurane gas mixed with pure oxygen and 150 μl of a cell suspension was injected into the right lower mammary fat pad (12 mice in each group; WT-Luc (control 1), Cas9-luc (control 2) and 4a-Luc (test group)). Mice were returned to their cages and allowed to recover. To analyze the luminescence from growing cells, mice were again anesthetized with 3 % isoflurane/O_2_ and then maintained at 2 % isoflurane/O_2_. Mice were then injected with a 200 μl mixture of saline with 15 mg/ml luciferin (Goldbio, St Louis, MO, USA) and the tumors imaged on an IVIS Lumina in vivo imaging system (PerkinElmer, Waltham, MA) every 30 s for up to 45 min. Luminescent images from peak levels (~20-30 min after injection) were analyzed using Living image 4.1 software (PerkinElmer) and ImageJ, calculating the integral and area of the luminescent signal above background that was recorded for each tumor. Mice wellbeing and tumor sizes were recorded in the mornings twice a week in the of procedure rooms in the Rush animal facility. Experiments continued until tumors reached 2 cm in size, until tumors reached 10% of body mass, or experience a moribund condition, whichever came first. The experiment was concluded after 5 weeks when 5 mice in the WT-LUC and CAS9-LUC met the requirements for euthanasia.

euthanized by CO_2_ asphyxia and the tumors resected and weighed immediately. Tumors were then frozen in RNAlater RNA stabilization solution (Millipore sigma, St Louis) and stored at −80 °C for subsequent use. Tumors were fixed in formalin and embedded in paraffin wax, sectioned and stained with hematoxylin/eosin and immunostained for Ki-67, the nuclear protein marker for cellular proliferation. Tumor section were photographed at x100 and x40 magnification from three fields of view of well stained areas. The level of staining was assessed by image deconvolution in ImageJ to assess the number of positively stained cells within each image. Statistical comparisons of tumor weight was performed by one-way ANOVA with significance assumed p<0.05.

### Seahorse XF analyzer measurements of extracellular acidification rates (ECAR)

WT MDA-MB-231 cells, Cas9 expressing cells, SRC, shRNA knockdowns and 2 knockout clones (4a and 5f_2_) cells grown in T75 tissue culture flasks were harvested at 70-90 % confluence and resuspended in DMEM 10% FBS, pen/strep at 100,000 cells/ml. 200 μl of this cell suspension was plated into a precalibrated Seahorse cartridge and allowed to attach overnight in 1% serum. Media was exchanged with Seahorse assay medium and the cells incubated in the absence of glucose for one hour. The cells were then analyzed per manufacturer instructions for real time extracellular acidification rate (ECAR) (Agilent Technologies, Santa Clara, CA. Briefly, the media was removed and replaced with Seahorse assay medium until they were placed in the Seahorse XF analysis machine. The Seahorse overlay had solutions of 1 M glucose, 10 mM oligomycin and 500 mM deoxyglucose (final concentrations of 10 mM glucose, 100 μM oligomycin and 5 mM deoxyglucose). The experiment was designed so that baseline recordings before addition of glucose were every 4 min, followed by readings every 8 minutes after glucose addition. Oligomycin was added 16 min after glucose addition and 2-deoxyglucose was added 16 min after the deoxyglucose. The units of mpH/min were calculated by the Agilent Seahorse XF analyzer software.

### Amplex Red assay of cellular hydrogen peroxide levels

Cells were grown as with the Seahorse measurements but were harvested on the day of the experiment and washed twice in ice cold Ringer’s solution without EGTA by centrifugation at 400 x g for 5 min. Cells were then resuspended in Ringer’s solution with 25 mM glucose added and kept on ice until the experiment. A stock solution of 100 μM Amplex red in Ringer’s solution was prepared and 50 μl of the reaction mixture was added to each well of a 96 well plate. 200,000 cells in Ringer’s suspension in 150 μl was added to each well and the release of H_2_O_2_ from the cells was recorded over the course of an hour in a Perkin Elmer Victor3V cell plate reader set at 37 °C with excitation wavelengths at 540 nM and an emission wavelength at 595 nM. A standard curve of H_2_O_2_ was performed with each experiment with concentrations of H_2_O_2_ running from 100 nM to 10 μM in order to calibrate the recording.

## Results

### Electrophysiological measurement of native H_V_1 in MDA MB 231 cells

The studies by Wang et al. [21, 22] showed expression of H_V_1 by western blotting and PCR but lacked functional data showing currents or pH changes. We examined MDA-MB-231 cells by patch clamp technique and recorded *bona fide* voltage-gated proton channels as determined by their kinetics, pH dependence, voltage dependence, proton selectivity, and sensitivity to Zn^2+^ (Fig 1). We were not always able to record definitive H_V_1 current in these cells, because extraneous conductance sometimes obscured the proton currents that were similar to conductance’s we reported previously in COS-7 cells [32]. In five out of nineteen cells proton currents were absent despite little obscuring noise. The cells varied quite considerably in size with capacitance ranging 7 – 92 pF with a mean value of 34.9 ± 25.4 (mean ± SEM, *n* =14). In 4 cells with definite proton current the proton conductance, *g*_H_, at pH 5.5/5.5 was 37.4 pS/pF. Fig 1A shows representative proton current families with 10 mV steps from −50 to 80 mV recorded from an MDA-MB-231 cell at 3 different pH gradients; pH 7/5.5, 6/5.5 and 5.5/5.5 (pHo/pHi). The current traces show clear pH dependent gating with currents becoming larger as the pH gradient increases from symmetrical pH to a 1.5-unit outward pH gradient. Currents at 80 mV in this cell were 27.5 pA at pH 5.5/5.5, 68.8 pA at 6/5.5 and 93.8 pA at pH 7/5.5. The reversal potential of the currents in this cell were 2.4 mV at 5.5/5.5, −24 mV at 6/5.5 and −66.5 mV at pH 7/5.5 and these values are plotted in Fig 1B in comparison with calculated *E*_H_ values as defined by the Nernst equation. Fig 1C shows conductance voltage curves for the representative cell with *V*_threshold_ values (defined as the voltage at which the *g*_H_ is 10% of its maximal value) of −28 mV at pH 7/5.5, 9 mV at pH 6/5.5 and 57 mV at pH 5.5/5.5. Currents reversed at −69 ± 14 mV at pH 7/5.5 (n=4) and at 8 ± 6 mV at pH 5.5/5.5 (n=4). Measured at 60 mV, the time constant for turn-on of current (τ_act_) was 3.2 ± 1.3 s at pH 7/5.5 and 6.3 ± 3.2 s at pH 5.5/5.5 (n=4). The current amplitude at 60 mV averaged 67.7 ± 38 pA (*n*=6) at pH 7/5.5 and 16.7 ± 18 pA (*n*=9) at pH 5.5/5.5. The currents were sensitive to Zn^2+^ as is characteristic of proton currents. Measured at pH_o_ 71 μM Zn^2+^ reduced currents and slowed the current turn-on in 3 cells.

**Fig 1.**
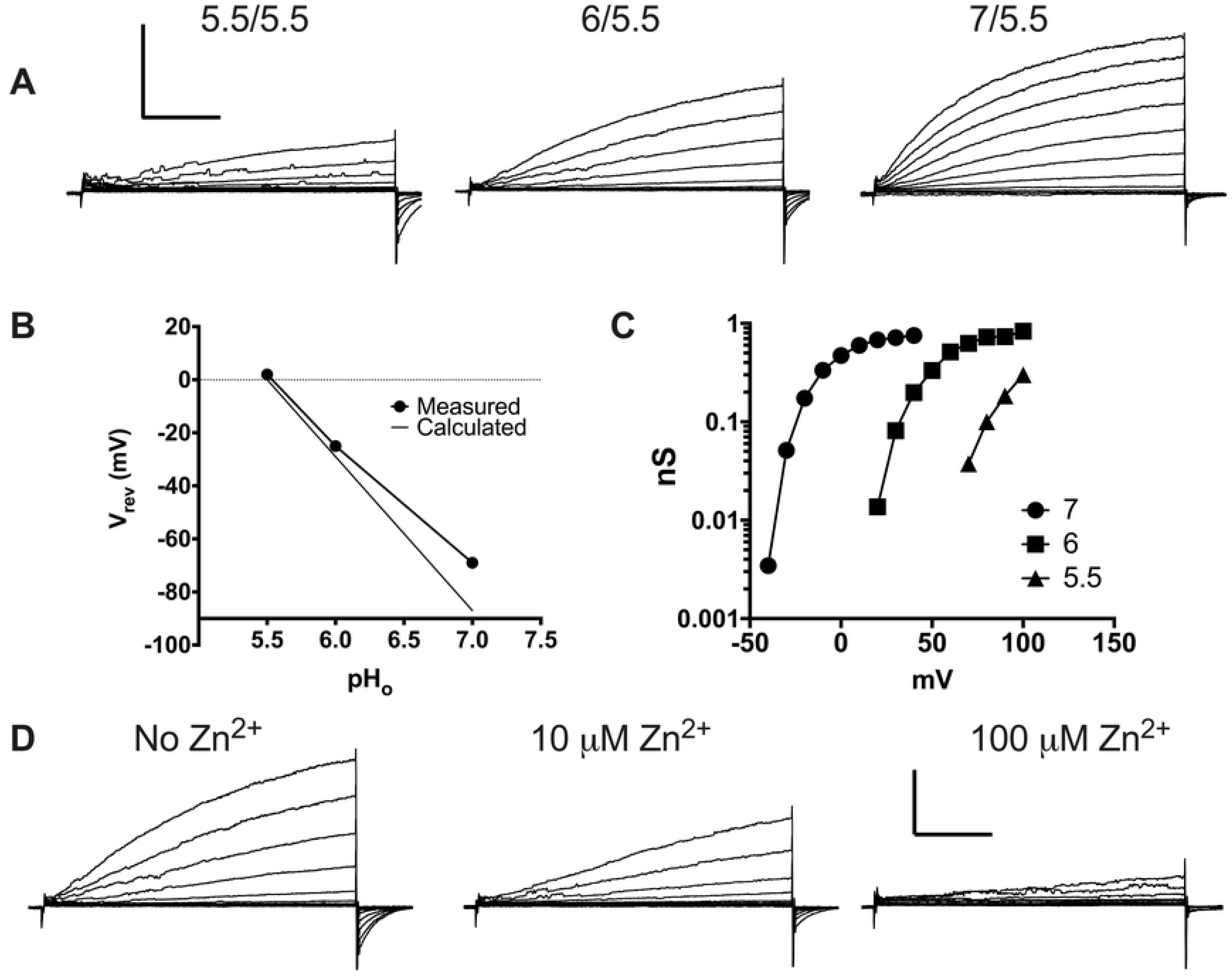
Proton current recorded in whole cell patch clamp configuration in MDA-MD-231 breast cancer cells. (**A**) Three families of currents at different pH_o_ all with a pH_i_ of 5.5 (labeled pH_o_/pH_i_). Voltage steps are every 10 mV. The scale bar units are 50 pA by 2 s. (**B**) The reversal potential (*V*_rev_) of a representative cell measured at each pH_o_ compared to the Nernst potential for protons, which assumes perfect selectivity for H^+^. (**C**) Proton conductance (*g*_H_) voltage curves derived from the data in **A**. (**D**) Three families of currents at pH 6/5.5 with different concentrations of Zn^2+^ as labelled. Voltage steps are 10 mV apart. The Scale bar units are 50 pA by 2 s.

### In-situ Staining of H_V_1 in Cells

Immunostaining of fixed non-permeabilized and permeabilized cells (Figs 2A and 1B, respectively) revealed strong staining in the plane of the membrane but little or no staining in the cytosolic plane of the cell when the cell was permeabilized to allow antibody access to the cell interior and H_V_1 cytosolic epitope, suggesting that the channel is expressed exclusively in the plasma membrane of the MDA-MB-231 cancer cell.

**Fig 2.**
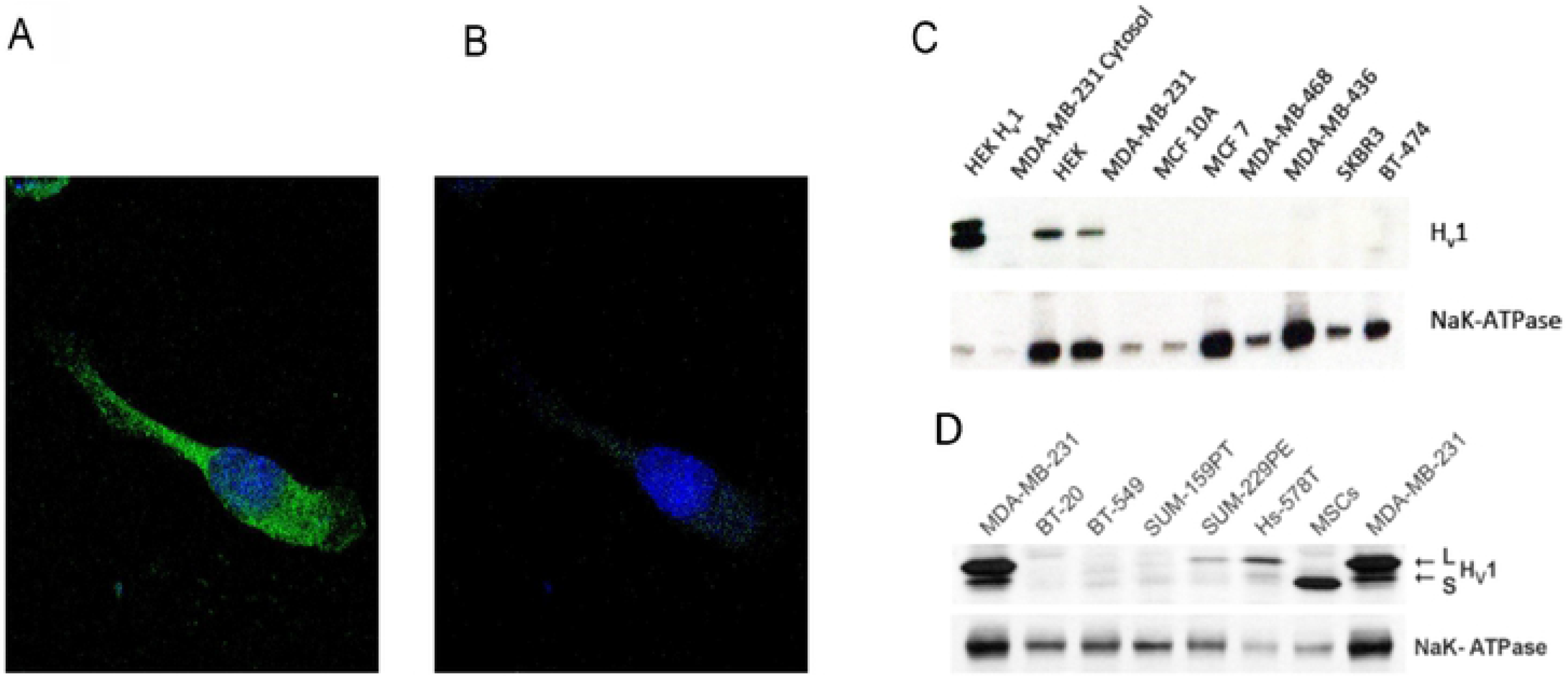
(A) A cell stained with a fluorescent antibody specific for H_V_1 with the image focused on the plane of the membrane, while **E** shows the fluorescence of the same cell but with the focus centered deeper into the cytosol. The nucleus was stained with DAPI and is visible in both **A** and **B**. (**C**) Western blot of membrane fractions from a number of different breast cancer cell lines. Membrane fractions were stained with antibodies specific to H_V_1 and Na^+^/K^+^ ATPase. The top pair of blots examines the membranes from a number of cells representative of the major types of breast cancer, i.e. Luminal A, Luminal B, Her2 and triple negative. (**D**) Western blots from a number of different triple negative breast cancer cells.

In order to find the optimal model system to study the effect of H_V_1 expression on cell function we screened 12 different breast cancer cell lines for H_V_1 using western blot techniques [33]. Cell types were chosen from a variety of different molecular classifications including luminal A, luminal B, basal-like, HER2-positive and normal subgroups. We homogenized cell membranes and examined the membrane lysates for the presence of H_V_1 protein (Figs 2C and 2D) and found evidence for significant expression of H_v_1 in 3 of them MDA-MB-231 cells, Hs578t and Sum229PE. Of these 3 cell lines MDA-MB-231 cells showed far higher expression than the other cell types. Attempts to record H_V_1 by patch clamp from HS-578t and Sum229PE cells were not successful. The cell lines that expressed H_V_1 are all classified as basal or triple negative breast cancer cell lines, because they do not express estrogen, progesterone receptors and the HER-2 protein. We chose to move forward with the MDA-MB-231 cells as a model because their H_V_1 expression was highest among the dozen cell lines tested.

### The effect of shRNA Knock-down (KD) of H_V_1 on the migration and H_2_O_2_ production of MDA-MB-231 cells

Wang et al. (2011) described a reduction in migration kinetics when using siRNA in the MDA-MB-231 cells to reduce the expression of H_V_1. We used a similar approach but with shRNA knockdown of H_V_1 in MDA-MB-231 cells. We tried several Mission shRNA sequences (Sigma-Aldrich, St Louis, MO) packaged in lentiviral vectors to transfect MDA MB 231 cells in order to reduce the expression of H_V_1 and found 2 shRNA sequences that were effective in knocking down the expression of H_V_1 in the MDA cells (Fig 3E). The successful sequences used, labeled 2.1 and 3.1 cells, reduced the expression of H_V_1 in MDA-MB-231 cells by 84 ± 1 % and 36 ± 4 % (mean ± SEM, *n*=3), respectively. We chose to replicate the migration assay by growing cells to confluence in 6-well plates, serum starving the cells for 24 hours and then wounding the monolayer with a sterile pipette tip. The cells were incubated in 0.5% serum medium to examine motility but not growth. Under these conditions the WT cells recovered 54.6 ± 7 % (*n*=4) of the wound in 24 hours whereas the cells transfected with the 2.1 and 3.1 shRNA constructs showed greatly reduced wound recovery of 11.6 ± 3% (*n*=4) and 13.3 ± 4% (*n*=4), respectively (Figs 3A and 3B).

**Fig 3.**
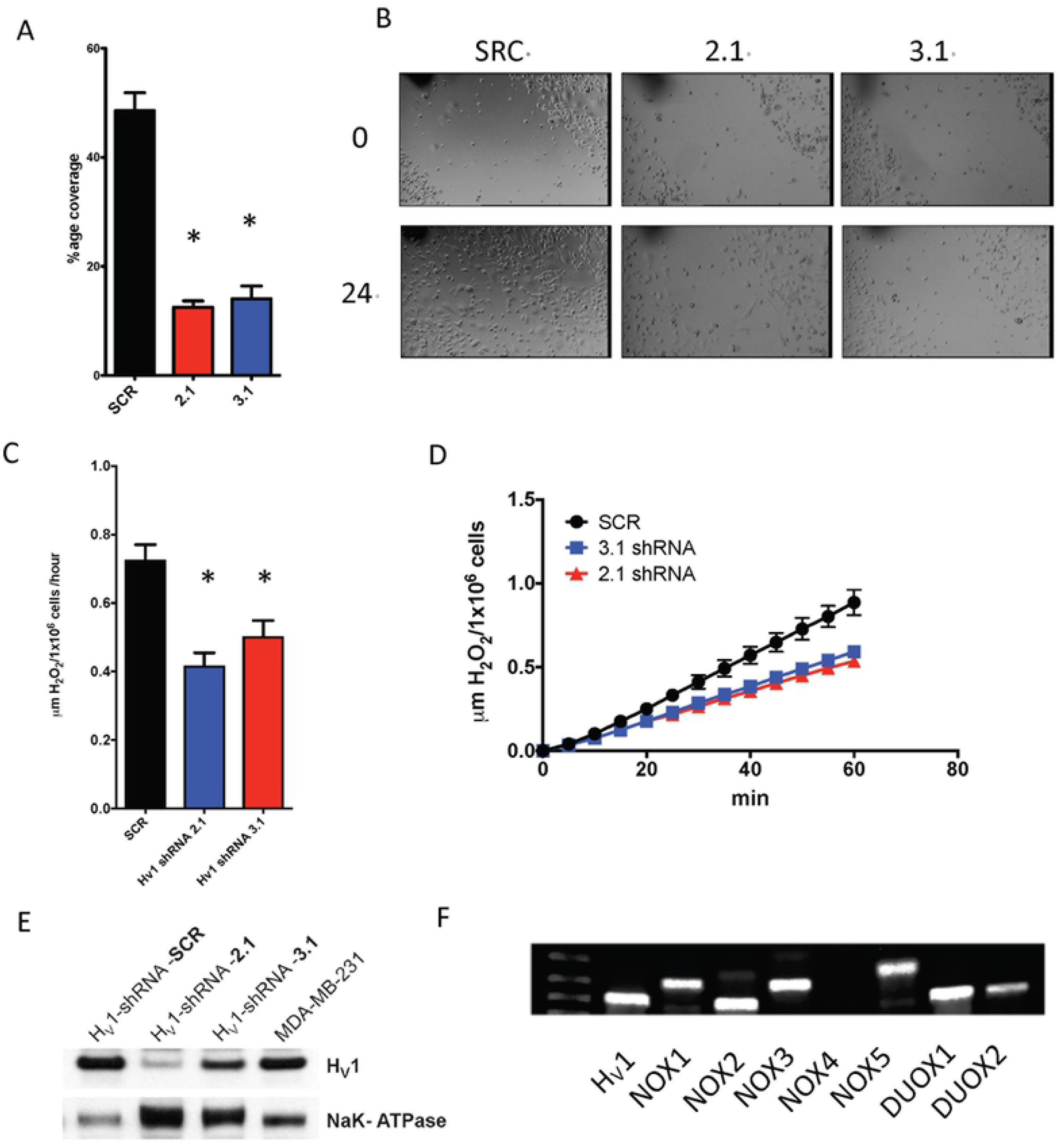
(**A**) Quantitative analysis of the wound healing assay shown in **B**. The chart shows the percentage coverage of each wound by cells after 24 hr. Analysis was performed using ImageJ and the chart shows data from 3 experiments performed in triplicate. Data are significant as determined by a normal one-way ANOVA with Bonferroni’s post-test *p*<0.0001. (**B**) Images of wells 24 hours apart after wounding with a 200 μl pipette. The black marks are scratch marks to denote position. MDA-MB-231 cells treated with virus containing the sequence for SCR, 2.1 or 3.1 shRNA for H_V_1 knockdown. Cells were kept in 1% FBS but with high glucose. (**C**) The release of H_2_O_2_ from MDA-MB-231 cells. The data is from 4 experiments performed in triplicate. 2.1 and 3.1 are significantly different from SRC as determined by a one-way ANOVA with Bonferroni’s post-test p<0.0015. (**D**) A representative experiment depicting the release of H_2_O_2_ over one hour. (**E**) A Western blot showing the inhibition of H_V_1 expression in MDA-MB-231 cells by shRNA 2.1 and 3.1. Na^+^/K^+^ ATPase was used as a loading control. (**F**) PCR amplification of products specific to all 7 members of the NOX family of membrane proteins.

H_V_1 is known to be closely associated with the activity of NADPH oxidases (NOX) in a number of cell types including neutrophils, eosinophils, B lymphocytes and basophils. Therefore, we test the hypothesis that inhibiting the expression of H_V_1 might result in a decrease in the production of superoxide, as has been demonstrated in numerous cell types. PCR analysis shows that MDA-MB-231 cells express the NADPH oxidase molecules NOX 1, 2, 3, 5 and both Dual oxidases (DUOX) isoforms (Fig 3F). We measured the release of H_2_O_2_ from cells treated with shRNA for H_V_1 or the scrambled control, SCR for an hour using Amplex red reagent. Cells transfected with shRNA specific to HVCN1 showed a significant reduction compared to SRC transfected control. Control cells released 0.72 ± 0.04 μmols H_2_O_2_/1×10^6^ cells/hour (*n*=6) compared to 0.42 ± 0.03 μmols H_2_O_2_/1×10^6^ cells/hour (*n*=4, *p*<0.001) for 2.1 shRNA and 0.5 ± 0.5 μmols H_2_O_2_/1×10^6^ cells/hour (*n*=4, *p*<0.01) for 3.1 shRNA.

### Construction of HVCN1 KO MDA-MB-231 cells via CRISPR gene editing

We started the shRNA experiments to replicate he work of others and generally corroborated previously published studies. However, some studies have shown that shRNA can have off target effects not seen with KO[34]. We were therefore deleted the functional HVCN1 gene altogether from these cells to examine how the cell functions in its absence. To this end, we stably transfected Cas9 into MDA-MB-231 cells (Fig 4A) before we targeted exon 2 of the H_V_1 sequence using the CRISPR guide sequence TTAAGGCACTTCACGGTCGT transfected into cells using a lentiviral vector. All splice variants of HVCN1 have the same initiating methionine; therefore, all splice variants were targeted [20]. We isolated several clones generated by this approach (Fig 4B). We were unable to detect expression of H_V_1 by western blot (Fig 4B) or the electrophysiological function of H_V_1 by the patch clamp technique (Fig 4C). Furthermore, sequence analysis of this clone showed that gene editing resulted in a 43-base insertion to one allele and an 83-base insertion into the second allele at the cut site for clone 4A. In clone 5f_2_, we observed in both cases a premature stop codon occurred shortly downstream of the cut site (Fig 4D).

**Fig 4.**
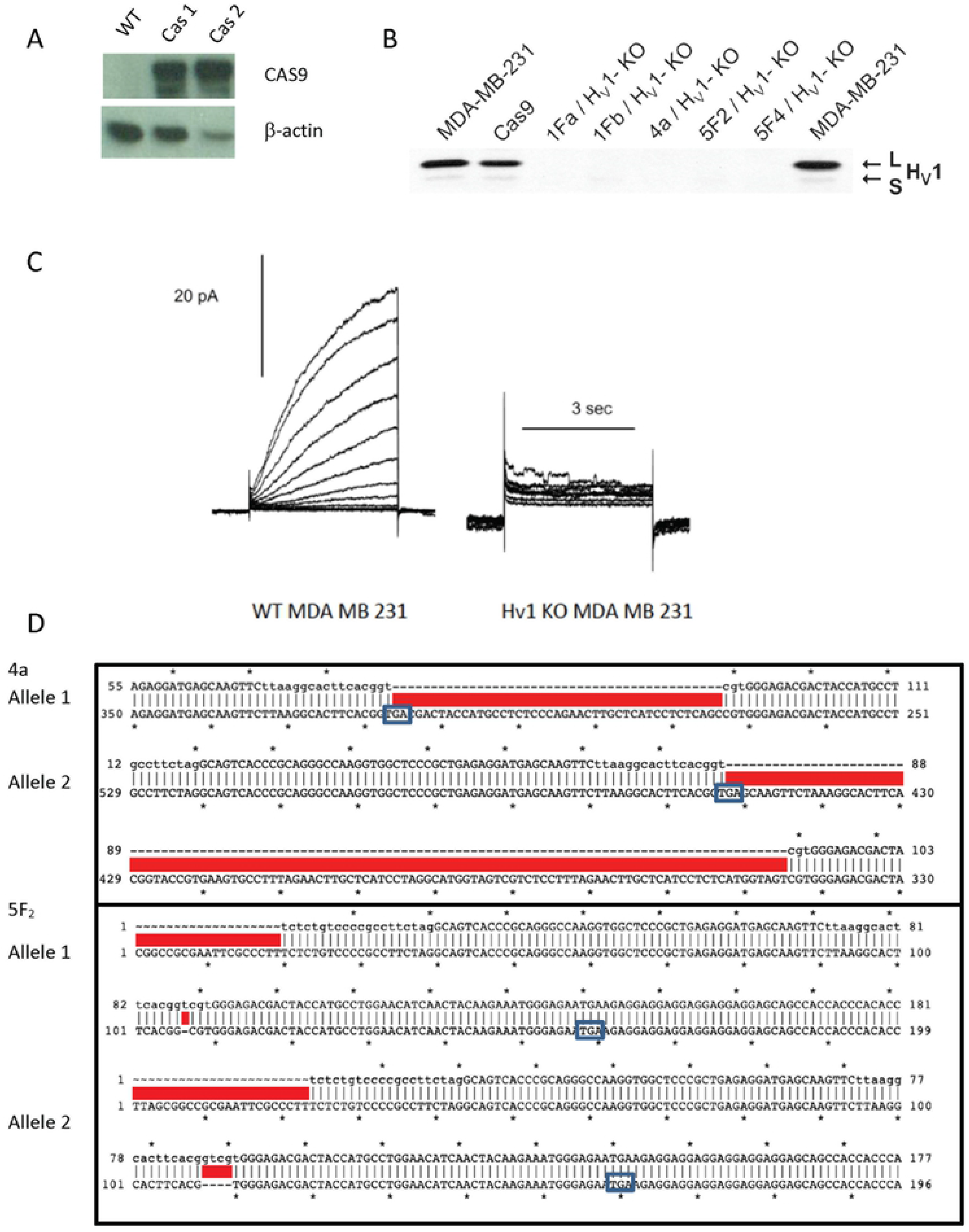
(**A**) Western blots specific to Cas9 showing the successful transduction of Cas9 protein in MDA-MB-231 cells. (**B**) A comparison of H_V_1 expression by Western blot in WT MDA-MB-231 cells and in 3 H_V_1 KO clones. (**C**) Comparison of current in WT MDA-MB-231 cells compared to a cell from H_V_1 KO MDA-MB-231 cells (4a). (**D**) The cut site of Cas9 on the 2 alleles for the HVCN1 gene in 4a (top) and 5f_2_ clones (bottom). Both alleles in the 4a clone were extended by nonhomologous end joining, the top allele by 43 nucleotides and the bottom by 83. In the 5f_2_ clone one allele had a 4 basepair deletion and one alelle had a single nucleotide deletion. In all cases a stop codon was created by frame shifts caused by the deletion or insert.

### The effect of H_V_1 KO on migration and H_2_O_2_ production of MDA-MB-231 cells

We compared the effect of knockout of H_V_1 to our results with knockdown cells by performing the same migration experiments on the 4a and 5f_2_ clones using the knock out cells, comparing the responses to Cas9 transfected cells. The Cas9 cells were chosen as the control because they have been manipulated in the same fashion, having been transfected with Cas9 and then clonally isolated before being transfected with null plasmid, whereas the knockouts were transfected with a guide sequence to the H_V_1 gene. The migration of the cells in low serum medium as measured by the percentage closure of a wound over 24 hours was 68.9 ± 5.8 % for the Cas9 control, 76.0 ± 6.4 % for 4a cells and 68.9 ± 5.2 % for the 5f_2_ cells (Fig 5A). None of these measurements were significantly different from each other as measured by ANOVA with Tukey’s multiple comparison posttest. When we measured the release of H_2_O_2_ from the different cell types (Fig 5C & 5D) we found that the KO cells produced more H_2_O_2_ than the Cas9-only expressing cells. The release of H_2_O_2_ from Cas9 cells was 0.23 ± 0.04 μM H_2_O_2_/1×10^6^ cells/hour (*n*=10) while the 4a cells released 1.52 ± 0.17 μM H_2_O_2_/1×10^6^ cells/hour (*n*=10) and the 5f_2_s release 0.85 ± 0.08 μM H_2_O_2_/1×10^6^ cells/hour (*n*=10). The release of H_2_O_2_ from the 4a cells was significantly different from the release of H_2_O_2_ from both the Cas9 cells and the 5f_2_ but neither the Cas9 cells nor the 5f_2_ were different from each other. This result seems to contradict the response observed for shRNA treated cells, although it should be noted that the Cas9 transfected cells released much less H_2_O_2_ than did SCR control MDA-MB-231 cells (Fig 3C).

**Fig 5.**
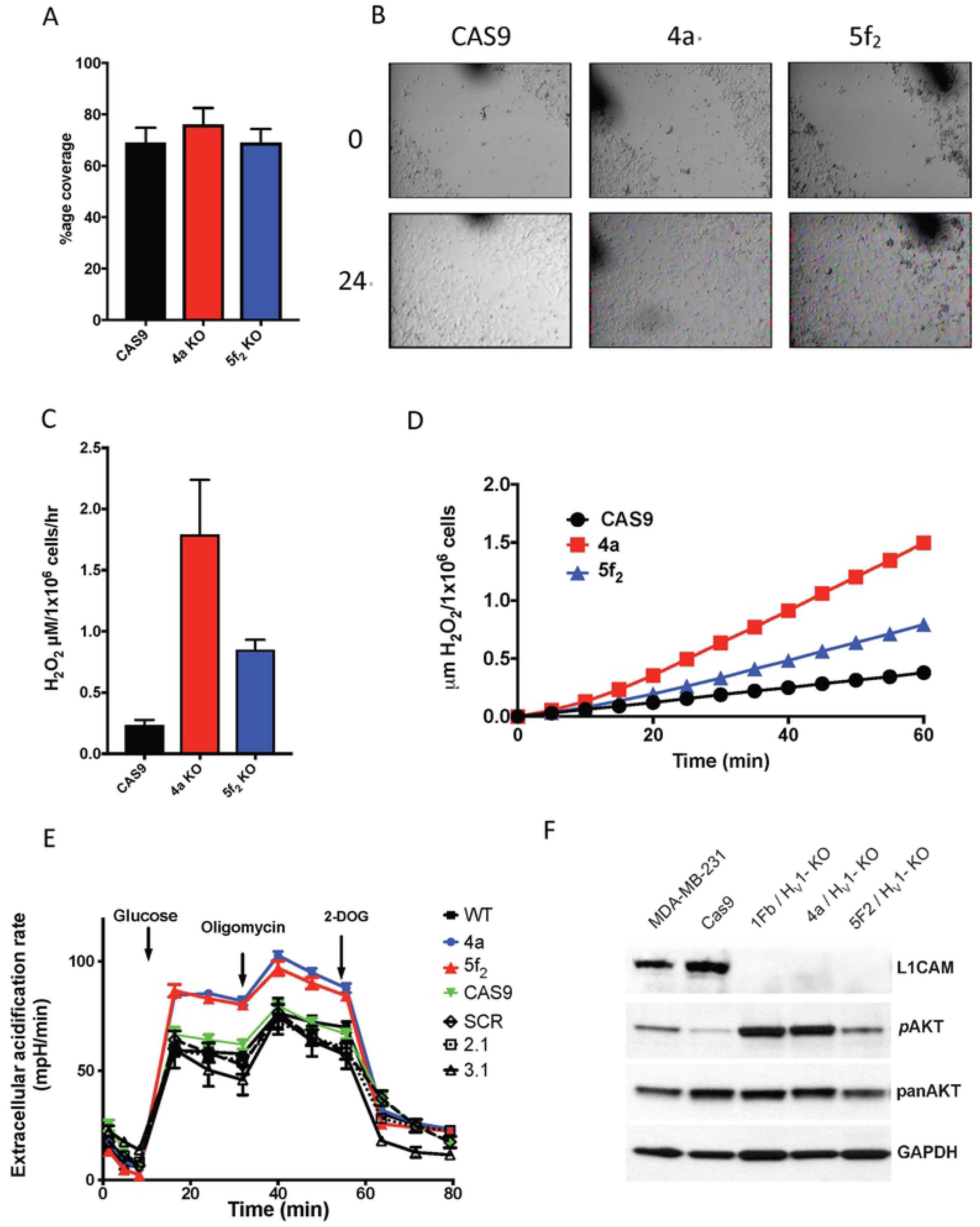
(**A**) Quantitative analysis of the wound healing assay shown in **B**. The chart shows the percentage coverage of each wound by cells after 24 hr. Analysis was performed using ImageJ and the chart shows data from 3 experiments performed in triplicate. Data are not significantly different. (**B**) Images of wells 24 hours apart after wounding with a 200 μl pipette. The black marks are scratch marks to denote position. MDA-MB-231 cells treated with virus containing the sequence for Cas9, 4a or 5f_2_ for H_V_1 knockout. Cells were kept in 1% FBS but with high glucose. (**C**) The release of H_2_O_2_ from MDA-MB-231 cells. The data is from 4 experiments performed in triplicate. 4a is significantly different from Cas9 as determined by a one-way ANOVA with Bonferroni’s post-test p<0.05. (**D**) A representative experiment illustrating the release of H_2_O_2_ over one hour. (**E**) The extracellular acidification rate of H_V_1 KO in 2 knockout clones compared to WT MDA-MB-231, Cas9, SRC, 2.1 and 3.1 knockdowns. Extracellular acidification was increased by adding 25 mM glucose after an hour of starvation. 10 μM oligomycin was used to inhibit mitochondria and enhance glycolysis. 2-deoxyglucose (5 mM) inhibits glycolysis to reveal the contribution of glycolysis to overall extracellular acidification. Data are the mean ± SEM from 5 experiments performed in at least sextuplet. (**F**) Western blot protein extracts from MDA-MB-231 cells showing the different intensities of protein expression or protein phosphorylation between WT and Cas9 expressing MDA-MB-231 cells and 3 different H_V_1 KO clones (1fb, 4a and 5f_2_).

As an additional comparison of the WT, Cas9 and KO cell lines, we analyzed the extracellular acidification rate (ECAR) of the various MDA-MB-231 cell types created. Fig 4E shows the result of starving the different cell types (WT, 4a, 5f_2_, Cas9, SCR, 2.1, 3.1) of glucose and then reinstating glucose and analyzing the extracellular acidification rate that follows. Under these conditions the H_V_1 KO cells showed a significant increase in ECAR with both 4a and 5f_2_ cells increasing to 84.3 ± 1.7 mpH/min and 86.6 ± 2.9 mpH/min (*n*=5) compared to 59.3 ± 2.7, 66.4 ± 3.3, 64.1 ± 4.1, 59.2 ± 2.6 and 59.8 ± 8.7 mpH/min for WT, Cas9, SRC, 2.1 and 3.1. The KO cells were significantly different from the other cells at *p*<0.001 after addition of glucose. Treatment of cells with oligomycin inhibits the mitochondrial electron transport chain and causes metabolism to shift completely to glycolysis. Treating the cells with oligomycin increased the ECAR values in all cells with the H_V_1 KO cells remaining significantly higher than the other cell types (*p*< 0.001) showing that they had a greater glycolytic reserve compared to non-KO cells.

Consistent with the enhanced ECAR values, we found an increase in phosphorylated AKT (Ser473) which is known to be a driver of glycolytic metabolism [35]. Assaying several different protein markers that were suggested from a reverse phase protein array screen, we found a reduction in the expression of the membrane adhesion molecule L1CAM to below detection levels in three different knockout clones (Fig 5F).

### Assessing the *in-vivo* growth of MDA-MB-231 WT cells compared to MDA-MB-231 H_V_1 KO cells

We next assessed whether the deletion of the functional H_V_1 gene would affect an *in-vivo* model of tumor growth. Previous studies [22] had shown that knockdown of the *HVCN1* gene resulted in slower tumor growth in a mouse model. Because we observed contradictory responses of knockdown and knockout cell models, we examined the growth of wild-type MDA-MB-231 cells compared to Cas9 expressing MDA-MB-231 cells and H_V_1 KO MDA-MB-231 cells. To monitor the growth of the tumors visually as well as via physical measurements we transfected the cells with a firefly luciferase plasmid in order to detect the growth of the tumor cells via luminescence without resorting to surgical methods. The cells were injected into the mammary fat pad of NSG/SCID mice and the mice were allowed free access to food and water for 36 days. By the end of the study the WT cells and Cas9 expressing MDA-MB-231 cells had grown significantly more than the 4a cells and had a mean weight of 537 ± 78 mg (*n*=12) and 753 ± 168 mg (*n*=10), respectively, compared to 197 ± 18 mg (*n*=10) for the H_V_1 KO cells (*p*<0.01). The physical appearance of the tumors after excision was also different. Tumors isolated from H_V_1 KO cells were small and white with a solid center. In contrast, the tumors from Cas9 and WT cells were larger and often necrotic and pus-filled in the center. In 3 animals from the Cas9 and WT groups, the tumors were not localized but appeared diffuse across the abdomen. The luminescence from these animals was also diffuse and showed signal all across the abdomen but these features were not seen in any animal from the KO group. Although there were differences in the luminescence patterns of the WT and Cas9 groups compared to the H_V_1 KO group, the total integral of luminescence of the WT, Cas9 and 4a groups did not significantly differ from each other.

## Discussion

Here we examined several possible mechanisms of action of H_V_1 in breast cancer cells using MDA-MB-231 cells as a model system. The inspiration of the study arises from studies by Wang et al., 2011, 2012 which suggested that proton channel expression in breast cancer cells was correlated with an increase in migration and a worse clinical outcome. Expression data in the METABRIC study [36] suggest that increased levels of H_V_1 expression occur in approximately 3% of breast cancer patients (58 out of 1905 patients) and is clustered almost entirely in the molecular subset of Claudin-low. Creation of Kaplan-Meier survival plots from data of patients in the METABRIC study who overexpressed H_V_1 did not show a significantly different survival rate compared to the general population of breast cancer sufferers, apparently in contradiction to the clinical data from the studies of the Wang et al., (Supplementary information 1). However, the interpretation of both sets of data are not straightforward, for the following reasons. The Wang et al., studies were based entirely on immuohistological data using an antibody that they had generated and has not been fully validated by them or others. However, we have evaluated 5 H_V_1 antibodies obtained from commercial sources with western blotting, and none of them convincingly detected human H_V_1 protein. In the present study we used an antibody generously provided by a collaborator (Dr. Melania Capasso, DZNE, Germany), that we validated in-house by overexpression of H_V_1 in HEK cells (Fig 2C) using a plasmid that we have used to generate proton currents previously [37]. It is conceivable that the Wang antibody results reflect non-specific immunoreactivity unrelated to the expression of the proton channel. Their published studies reported no thorough attempt at validation of the antibody at a basic level. Conversely, the METABRIC study examines only mRNA at the cellular level; it is well-established that mRNA and protein levels are can be poorly correlated [38, 39]. Therefore, even though there is good evidence that HVCN1 mRNA was upregulated in the 58 patients from that study, no actual protein expression data exist to corroborate the mRNA expression.

Our data indicate that three breast cancer cell lines express readily detectable H_V_1 via western blotting, namely sum229PE, Hs578t, and MDA-MB-231. However due to the low intensity of antibody staining and the absence of convincing patch clamp recording of proton current we cannot be sure that the first two of these cells express sufficient levels of H_V_1 to significantly affect cell function. We chose to focus on MDA-MB-231 cells because H_V_1 expression was highest in this cell line and proton current was confirmed electrophysiologically. Nevertheless, some MDA-MB-231 cells appeared to lack measurable H_V_1 current. Whether some cells in this cell line do not express H_V_1 at all or whether the expression level is low enough in some cells to prevent detection of current is unclear. In any case, the expression of H_V_1 in this cell line is heterogeneous, which might suggest the existence of different phenotypes within this cell line, a phenomenon previously noted [40], thus the possibility that only a subset of these cells express H_V_1 cannot be excluded [41]. Certainly, most of the MDA-MB-231 cells recorded here expressed *bona-fide* H_V_1. Unequivocal proton currents were measurable by the patch clamp technique and their properties, including opening and closing kinetics, pH sensitivity, selectivity, and inhibition by Zn^2+^, all resemble analogous properties of H_V_1 in other cells (alveolar epithelial cells, neutrophils, eosinophils, macrophages, monocytes, sperm, oocytes, etc).

An Examination of the Reactive Oxygen Species Hypothesis Our initial experiments repeated and extended data from the Wang et al. studies. We evaluated changes in motility of the cells and to this end we used shRNA inhibitory sequences derived from the mission RNA database. We tried 5 sequences of which 2 were effective in reducing H_V_1 expression to a significant extent. Our wound scratch test experiment confirmed that knockdown of H_V_1 inhibited the movement of cells over 24 hours (Figs 5A & 5B), consistent with the previous study. We postulated that changes in NADPH oxidase activity could be a plausible mechanism for the action of H_V_1 in these cells, because of close relationship between NADPH oxidase and H_V_1 in neutrophils, eosinophils, B-cells and monocytes where its role is well studied and established [15, 42–44]. We measured mRNA of NADPH oxidase variants, including Nox1, Nox2, Nox3, Nox5, Duox1 and Duox2, in these cells, which established that the molecular machinery necessary for producing reactive oxygen species were present. Measuring the release of H_2_O_2_ from knockdown cells compared to a scrambled shRNA plasmid showed a significant reduction in the release of H_2_O_2_ although the release levels appeared to correlate with the level of H_V_1 expression only qualitatively. The cells transfected with the 2.1 plasmid did not show a significantly greater reduction in the release of H_2_O_2_ compared to those transfected with 3.1 plasmid even though the differences in H_V_1 protein expression are 21% and 65% of wild type respectively (Fig 3E).

### Comparison of knockdown (shRNA) and knockout (CRISPR/Cas9) of HVCN1

One might expect that functions inhibited by the knockdown of H_V_1 might show a greater level of inhibition if the channel were removed completely. Therefore, we created knockout cells using CRISPR/Cas9 to delete the HVCN1 gene. We targeted a sequence at the beginning of exon 2 of the gene to ensure that we removed expression of both long form and short form variants [20]. To achieve this, we first transfected Cas9 stably into MDA-MB-231 cells using Blasticidin as a selection tool and clonally isolated a cell from the selection. Using this clonal line, we targeted the gene using a guide sequence that created several clones that were absent in the expression and function of the proton channel as assessed by patch clamp and western blot and sequence showed that non-homologous end joining created a cut in the site specified. Thus, all knockout cells came from the same clonal cell and the Cas9 cells served as an additional control to wild type. Of the knockout cells isolated many died shortly after creation with the clones 5f_2_ and 4a being the most stable; however only the 4a KO cells survived long enough to withstand labeling with luciferase and incorporation into a mouse xenograft model. When we measured migration and H_2_O_2_ release from the knockout cells compared to the Cas9 control we did not observe inhibition of either migration or H_2_O_2_ release, in contrast to the results seen with shRNA knock-down of H_V_1. Compared to WT MDA-MB-231 cells, H_2_O_2_ release was slightly lower, but Cas9 transfected cells generally had much lower H_2_O_2_ release that WT (2.08 ± 0.5 μmols H_2_O_2_/1×10^6^ cells/hour, N=10). Migration of the Cas9 cells was faster than WT cells and this phenotype was maintained in the KO cells where migration was not significantly different among Cas9, 4a and 5f_2_ cells (Fig 5A).

We employed a reverse phase protein array available through MD Anderson Cancer Center (University of Texas; Houston, Texas) to screen differences in protein expression and phosphorylation of ~300 targets in the KO cells compared to WT and Cas9 (Supplementary Figure 2). LlCAM, an adhesion molecule that is associated with poor prognosis in certain breast cancers [45, 46], was expressed at much lower levels in the KO cells. We also detected a higher level of AKT phosphorylation (Ser473) in the KO cells. There was functional correlation with this increase in phospho-AKT as the KO cells we analyzed for ECAR had higher output than the WT, Cas9 and shRNA cells, regardless of whether the mitochondria were inhibited.

The evidence we accumulated showed that knockdown and knockout cells functioned in qualitatively different fashions. Given that Wang et al. [21, 22] found that inhibiting the expression of H_V_1 with shRNA slowed tumor growth, we wondered whether the complete KO of H_V_1 from MDA-MB-231 cells would do the same. We chose WT and Cas9 cells as controls because the Cas9 had functional differences compared to the WT (smaller H_2_O_2_ release, faster migration) and the Cas9 cells were the parental clones of the KO cells. The intent was to minimize the changes attributable to the genetic modification. In agreement with previous data, the KO cells developed into significantly smaller tumors over 35 days compared to both WT and Cas9 cells. We had labeled the cells with a luciferase enzyme that allowed us to monitor changes in luminescence via a bioanalyzer for the duration of the experiment. Taking the integral of luminescence, we found no significant difference between luminescence of the control compared to KO cells. The reasons for this are not clear but may reflect the manner in which luminescence is generated. The luciferase reaction requires oxygen and thus changes in the perfusion of the tumors by blood can alter the signal. The smaller tumors of the KO group may have been better perfused than the WT and Cas9 cells, the latter resulting in a large tumor but with lower luminescence. Another possibility is that the overall number of tumor cells remained the same, but the WT and Cas9 cells tended to migrate away from their initial location. The resulting damage and necrosis in the inner regions of the tumor may have resulting in a necrotic tumor center with an outer layer of tumor cells that were similar in number compared to the KO tumors. In four animals injected with WT or Cas9 tumors the luminescence spread across the abdomen (Fig 6A) and in one case migrated from the injected side to the opposite mammary gland. There was the appearance of greater mobility of the control group luminescence compared to the very predictable size and location of luminescence of the KO tumors.

**Fig 6.**
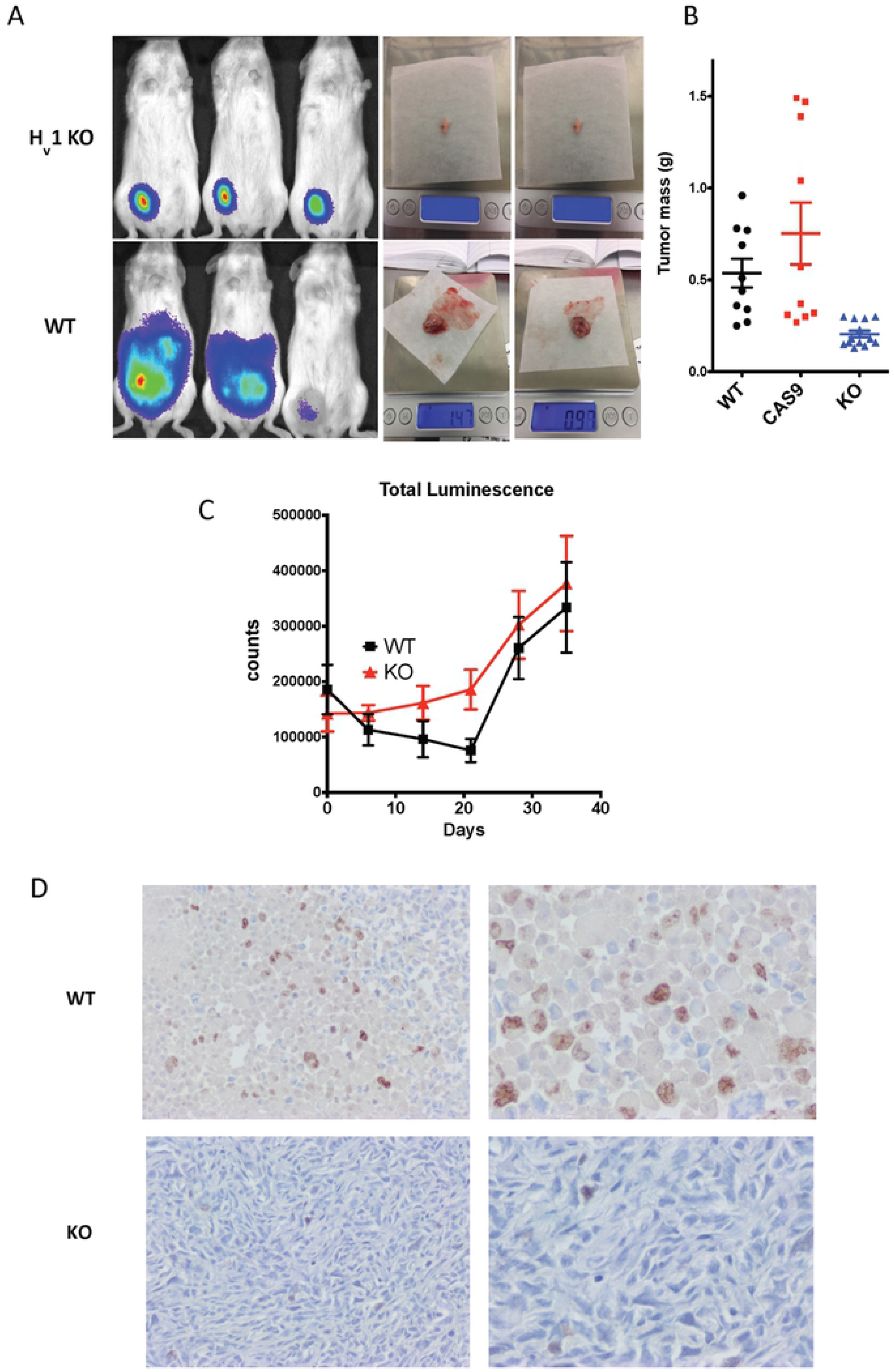
(**A**) Three representative mice inoculated with tumors in their mammary fat gland at the end of 35 days of the experiment. Top left panel shows mice with tumors from H_V_1 KO 4a cells and the lower left panel shows tumors from wild type cells. The center panels shows 2 representative tumors from each group on weighing scale shortly after dissection and blotting for fluids. (**B**) A dot plot graph that categorizes the different weights of the various tumors that were resected from the mice in each group. The mean ± SEM are plotted in each case. There were significant differences between the 4a group and the WT and Cas9 group (*p*<0.05). (**C**) A graph comparing the integrated luminescence intensities of tumors as measured every seven days by injection of 200 mg/kg luminol into each mouse and allowing for the luminescent intensity to plateau. The luminescence counts are normalized to the relative expression determined by luminescence intensity measured in 1×10^6^ cells in-vitro. (**D**) shows 4 images at x40 (left) and x100 (right) of 2 representative tumors taken from the WT group (top) and 4a group (bottom) that were stained for Ki67 (brown staining).

In attempting to understand how H_V_1 might alter tumor growth, the decreases in H_2_O_2_ production and migration seen in the shRNA cells may not be causally relevant, because the KO cells showed neither of these traits yet exhibited significantly smaller tumor growth. Similarly, the significance of the increase in ECAR is complicated by the fact that our shRNA cells did not show the same phenotype yet siRNA treated cells in the Wang et al. study also exhibited reduced tumor growth.

The resulting locally “inverted” acidic cellular environment seen in acidic tumors been proposed to influence the invasive potential of tumor cells via the acid-mediated invasion hypothesis [45]. Gatenby and Gawinski proposed this hypothesis in 1996, describing the process of acid-mediated invasion as increased acid production serving as an intermediate by causing H^+^ flow down concentration gradients into adjacent normal tissue [47]. The chronic exposure of surrounding healthy tissue to an acidic microenvironment produces toxicity and cell death by the collapse of the transmembrane H^+^ gradient inducing necrosis or apoptosis [48]. The increased extracellular acidification enhances extracellular matrix degradation through the release of cathepsin B and other proteolytic enzymes. In contrast, tumor cells, due to their enhanced pH regulatory mechanisms, are resistant to the acid-induced toxicity, allowing them to survive and proliferate in the low pH microenvironments [49]. This permits the tumor cells to invade the altered adjacent normal tissue despite the acid gradients. In vivo experiments and modeling experiments have shown evidence for the presence of peritumoral acid gradients as well as cellular toxicity and extracellular matrix degradation in the normal tissue exposed to the acidic microenvironment [50]. Further evidence supporting this theory includes observations that neutralization of the tumor-derived acid with systemic buffers (bicarbonate) was sufficient to inhibit spontaneous and experimental metastases [51].

The presence of H_V_1 in late stage breast cancer at first glance appears to be in line with the upregulation of other pH transporters seen in cancer tissue. However, the mechanism of action of H_V_1 runs contrary to the development of an inverse proton gradient that precedes the invasion as predicted by the model of the acid-invasion hypothesis. Under the conditions of an inverted pH gradient the proton channel would be closed and therefore would be unlikely to affect tumor growth. With the evidence suggesting that H_V_1 does affect tumor growth then it must occur in microenvironment circumstances that differ from the acid-invasion hypothesis because the only way the channel would open and function is if the cell were exposed to an outward pH gradient during its growth cycle. The evidence suggesting that H_V_1 enhances tumor growth and metastasis in breast cancer suggests that the acid-invasion hypothesis is not a complete picture of the factors that contribute to cellular invasion and tumor growth.

RPPA analysis and RNAseq analysis (Supplementary Figure 3) of the KO cells compared to WT and Cas9 reveal a number of genes with altered expression; including downregulation of L1CAM, ECAM, b-catenin and upregulation of AKT2 and AKT-p473. It is tempting to speculate on the mechanism of H_V_1 based on how the proton channel works (it opens when the proton electrochemical gradient is outward and closes when the gradient is inwards). The changes in protein expression in the absence of H_V_1 (downregulation of adhesion molecules and upregulation of metabolism) suggests that the cell uses H_V_1 as a sensor of its external pH environment. In conditions that result in H_V_1 opening, the increase in cell adhesion molecules would facilitate moving through the microenvironment; whereas when H_V_1 closes the cell increases its metabolism to enhance survival in an externally acidic environment. This idea is consistent with the observation that tumors grow slower in the absence of H_V_1.

In summary, we show further evidence that H_V_1 contributes to the growth of tumors created by the injection of the triple negative breast cancer cell line MDA-MB-231 into the mammary fat pad of mice and provide evidence that H_V_1 expression modulates aspects of metabolism and the expression of adhesion molecules.

## Acknowledgements

This work was supported by National Institutes of Health Grants GM121462 and GM126902 (to T.E.D.), Bears Care and The Brian Piccolo Cancer Research Fund (DM and AMA).

